# Kin-recognition shapes collective behaviors in the cannibalistic nematode *Pristionchus pacificus*

**DOI:** 10.1101/2023.07.14.549064

**Authors:** Fumie Hiramatsu, James W. Lightfoot

**Affiliations:** Max Planck Research Group Genetics of Behavior, Max Planck Institute for Neurobiology of Behavior – caesar, Ludwig-Erhard-Allee 2, 53175, Bonn, Germany

**Author notes:** Corresponding author at.

## Abstract

Kin-recognition is observed across diverse species forming an important behavioral adaptation influencing organismal interactions. In most species, proximate level mechanisms are poorly characterized, but in the nematode *Pristionchus pacificus* molecular components regulating its kin-recognition system have been identified which determine its predatory behaviors. This ability prevents the killing of kin however, its impact on other interactions including collective behaviors is unknown. Utilizing pairwise aggregation assays between distinct strains of *P. pacificus*, we observed aggregation between kin but not distantly related con-specifics. In these assays, only one strain aggregates with solitary behavior induced in the rival. Abolishing predation through *Ppa-nhr-40* mutations results in rival strains successfully aggregating together. Additionally, interactions between *P. pacificus* populations and *Caenorhabditis elegans* are dominated by *P. pacificus* which also disrupts *C. elegans* aggregation dynamics. Thus, aggregating strains of *P. pacificus* preferentially group with kin, revealing competition and nepotism as previously unknown components influencing collective behaviors in nematodes.

## Introduction

Kin-recognition is observed across many species and is associated with a diversity of different behaviors. This includes behaviors in single celled organisms such as swarming behaviors in bacteria (*1*), flocculation in yeast (*2*), and altruism in the amoeba *Dictyostelium* (*3–5*). Additionally, behaviors in more complex organisms are also thought to be reliant on kin-recognition including cooperation in insects (*6*, *7*) and lizards (*8*, *9*), colony fusions events in tunicates (*10*, *11*), cannibalism in several amphibian species (*12*, *13*), as well as nest mate preference in rodents (*14*, *15*). In nematodes a kin-recognition system was also recently identified in *Pristionchus pacificus*. This is an omnivorous and cannibalistic nematode species and is capable of attacking and killing the larvae of other nematodes as well as other *P. pacificus* con-specifics (*16*, *17*). These predatory behaviors likely provide a supplementary nutrient source as well as a mechanism to remove potential competitors from their environment as both territorial behaviors and surplus killing have been described (*18*–*20*). Furthermore, while *P. pacificus* can be a voracious predator of other nematode larvae, it avoids killing its close relatives and progeny through the existence of a hypervariable small peptide mediated kin-recognition system (*21*, *22*). However, previous *P. pacificus* kin-recognition studies have focused on the interactions between predators and larvae, and little is known of its impact on larger scale group behaviors. Importantly, these larger scale dynamics are likely to occur frequently due to the boom-and-bust life history strategy employed by *P. pacificus* in its natural ecological setting which temporarily results in a large concentration of nematodes around a confined niche (*23*).

While few studies on larger scale collective dynamics have taken place in *P. pacificus*, aggregation behaviors have previously been reported in a high-altitude adapted clade as a mechanism to avoid hyperoxia although it is likely other as yet unstudied factors will also induce aggregation in this species (*24–27*). Instead, nematode aggregation behaviors have been intensely studied at the molecular and neuronal level in the model nematode *Caenorhabditis elegans.* These behaviors in *C. elegans* are induced in response to various factors including hyperoxia, aversive stimuli, food quantity and population density (*28–30*). Accordingly, many wild isolates of *C. elegans* show aggregation behaviors in which they group together on a bacterial lawn and even exhibit dynamic swarming over longer time periods (*31*, *32*). In contrast, the *C. elegans* laboratory reference strain N2 does not aggregate due to the presence of a gain-of-function mutation in the *npr-1* gene and is considered to be a solitary strain (*28*, *33*). This behavioral difference is caused by a single amino acid change in this neuropeptide receptor with gregarious wild isolates carrying a 215F allele version of *npr-1* while in solitary strains a 215V version is found instead which suppresses this behavior. However, both variants have been shown to confer fitness advantages depending on food availability and dispersal strategy (*34*, *35*). In addition to *npr-1*, aggregation is mediated through various receptors found in sensory neurons which depend on the intraflagellar transport (IFT) machinery, several soluble guanylate cyclases and components of the transforming growth factor beta (TGF-β) family (*29*, *30*, *36*, *37*). This includes the TGF-β *daf-7* pathway which acts in parallel to *npr-1.* Furthermore, pheromone signaling through small molecule ascarosides also promotes attraction and aggregation which converge on the same neuronal circuits (*38–40*). Intriguingly, while the IFT system is essential for aggregation in *P. pacificus*, neither *Ppa-npr-1* nor *Ppa-daf-7* are required for these behaviors (*26*, *27*, *41*). Therefore, aggregation in *P. pacificus* is dependent on a distinct mechanistic process that has evolved independently from those described in *C. elegans*.

Here, by exploring the specialized high-altitude adapted and aggregating clade of *P. pacificus* we identify kin-recognition as an essential component shaping aggregate formation. We find aggregating strains of *P. pacificus* preferentially group with their own kin or close relatives and avoid more divergent strains. Moreover, pairwise interactions between distantly related strains reveal that one strain frequently dominates and aggregates while the other is displaced. This territoriality also extends to interactions of *P. pacificus* with other aggregating species including *C. elegans* and reveals kin-recognition and competition as a previously unknown component influencing collective behaviors in nematodes.

## Results

Collective behaviors are found in many nematode species indicating its high degree of evolutionary conservation (*42*). Conversely, In *P. pacificus* most strains behave in a solitary manner with aggregation only predominantly observed in one of the main *P. pacificus* evolutionary lineages referred to as clade B. However, it is also likely that other unexplored factor may also induce aggregation in this species more generally (Fig. 1A-C). Strains belonging to this clade are frequently found in a high-altitude environment and have adapted to lower oxygen concentrations. As such, under normal laboratory conditions, aggregate formation is induced and animals localize at the border of the bacterial lawn to avoid hyperoxia (*24–27*). Furthermore, unlike in *C. elegans* this aggregation behavior is observed even in the absence of a bacterial food source (Fig. S1). Importantly, like all other strains of *P. pacificus* sampled so far, isolates from these locations are also predatory and kill the larvae of other nematodes including other *P. pacificus* con-specifics (Fig. 1D). They also possess kin-recognition behaviors which prevent the killing of their own progeny and close relatives while enabling the killing of more distantly related strains (*21*, *22*). Therefore, we exploited the *P. pacificus* clade B propensity to aggregate to investigate the influence of kin-recognition and its associated predation on group behaviors in these nematodes.

**Figure 1.**
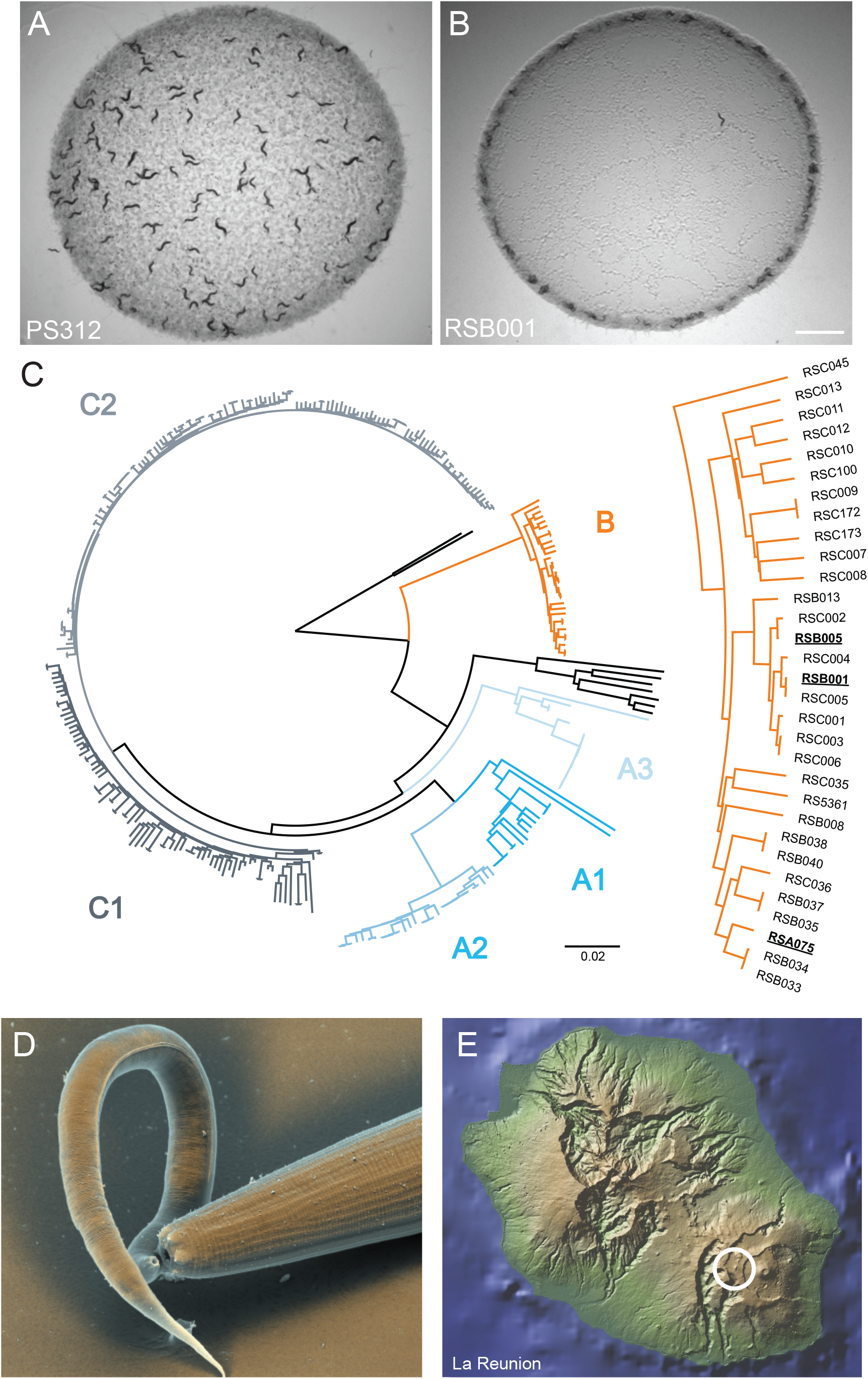
*P. pacificus* aggregation behaviors are prevalent in a high altitude adapted clade. (A) The main laboratory *P. pacificus* wild type strain, PS312 does not aggregate under standard laboratory conditions. (B) RSB001 is a high altitude adapted strain which aggregates under standard laboratory conditions. Scale bar = 2000 µm. (C) *P. pacificus* phylogenetic tree representing the genetic relationship between 323 wild isolates. Image adapted from Rödelsperger et al 2017 (*44*). The three strains selected for further analysis are highlighted in bold. (D) SEM image of a *P. pacificus* predator (large right nematode) killing a *C. elegans* larvae (smaller left nematode). (E) Strains selected for analysis were all isolated from the region circled which is a high-altitude environment on La Reunion Island.

Accordingly, we selected three strains from one high altitude location to investigate their interactions and ability to form aggregates together. These strains were all originally isolated from the same area on the Nez de Bœuf volcano peak (>2089 m above sea level) on La Reunion Island (Fig. 1E) and as such also represent potentially ecologically relevant interactions. We selected RSB001 as it has previously formed the basis of numerous molecular studies (*25*, *27*, *43*, *44*) and two strains of increasing phylogenetic distance, the relatively closely related strain RSB005, and a more divergent strain, RSA075 (Fig. 1C).

### Kin-recognition and predation influence aggregation behaviors

Having identified aggregating strains with differing degrees of genetic relatedness, we next explored if aggregation behaviors occurred between these mixed *P. pacificus* populations. To assess this, we established aggregation assays in which 120 worms (60 worms from each strain) were placed onto a defined OP50 *Escherichia coli* bacterial lawn of 70 µl for 3 hrs. In order to differentiate and define strain specific behavioral interactions in mixed populations, we utilized a previously established staining method using fluorescent vital dyes that has been demonstrated to have no effect on the animal health (*45*) or its aggregation behavior (Fig. 2A and fig. S2A). Under these conditions, all three strains aggregated strongly in control assays consisting of only their own strain (Fig. 2B-C and fig. S2B). However, in mixed assays consisting of two strains, we observed striking differences in aggregation depending on the combination of strains assessed. Specifically, aggregates between populations of the relatively closely related RSB001 and RSB005 were consistent with controls and aggregated together abundantly. However, assays between either RSB001 or RSB005 together with the more distantly related RSA075, resulted in a significant decrease in aggregate formation with solitary behaviors becoming more prevalent (Fig. 2B and 2C and Movie S1). Furthermore, the composition of aggregates also changed in mixed populations between more distantly related strains. While aggregation was disrupted for both strains, we observed that one strain, RSA075, was able to dominate and formed more aggregates than both RSB001 and RSB005 during pairwise interactions (Fig. 2B to D).

**Figure 2.**
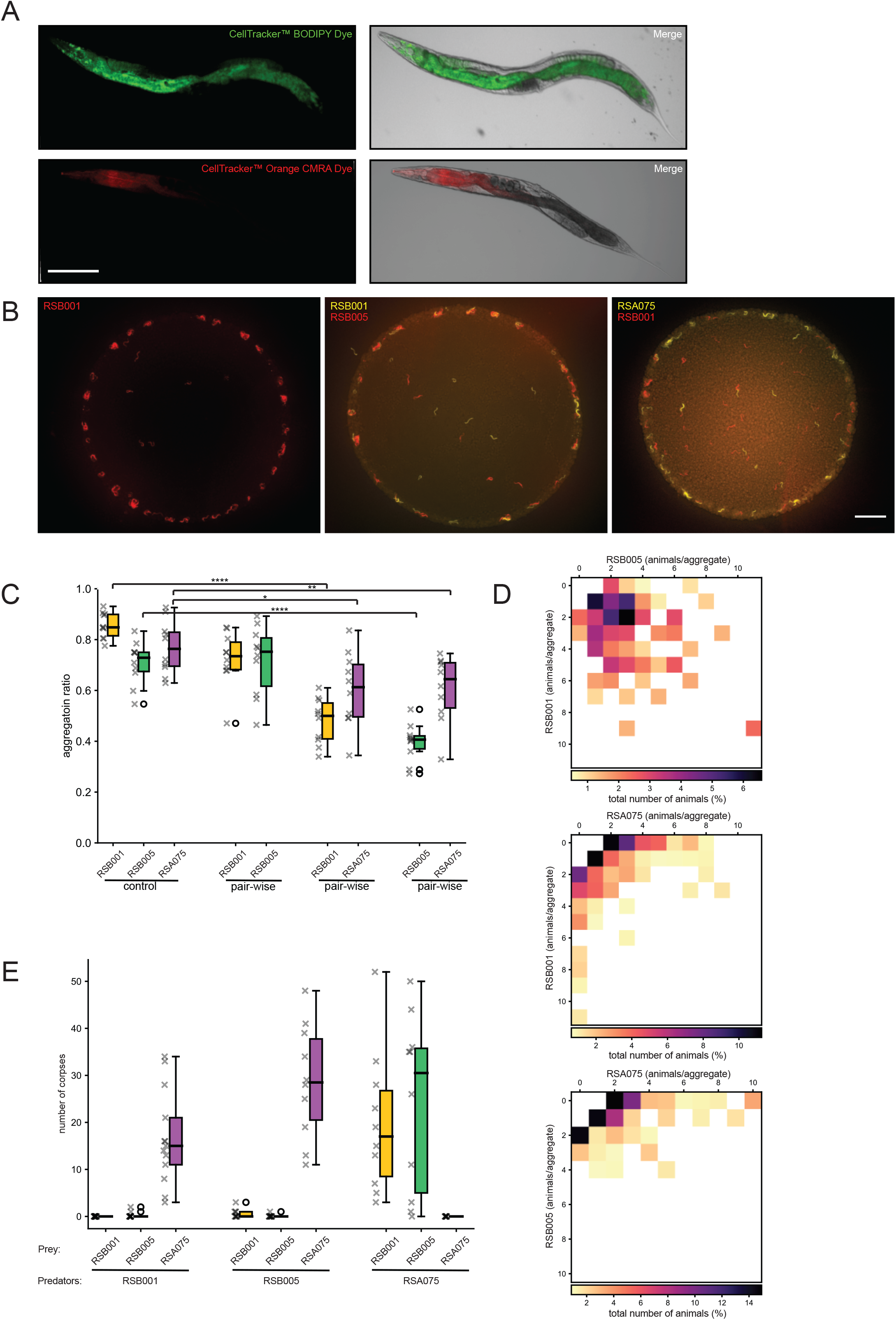
***P. pacificus* aggregates with kin and displaces non-kin.** (A) Vital dyes allow fluorescent labelling of distinct populations for identification in mixed cultures. Scale bar = 200 µm. (B) Representative images of the aggregation phenotypes of the *P. pacificus* strain RSB001 control as well as mixed cultures of RSB001 with RSB005 and RSB001 with RSB075. Scale bar = 2000 µm. (C) Quantification of aggregation ratios of *P. pacificus* cultures including RSB001, RSB005, and RSA075 controls as well as all pairwise permutations. Aggregation is disrupted between more distantly related strains. n = 10 per condition. p<0.0001 for RSB001 and RSB005 with RSA075, p<0.05 for RSA075 with RSB001 and p<0.01 for RSA075 with RSB005. (D) Heatmaps of pairwise interactions revealing the proportion of each strain in an aggregate. Between strain aggregates are frequent among RSB001 and RSB005 but monotypic aggregates are more prevalent when either strain is paired with RSA075 which also dominates the aggregate formation (E) Quantification of predation assays revealing killing between all possible pairwise permutations. Increased killing is observed between more distantly related strains. n = 21 for RSB001 predators on RSB001 prey, n = 15 for RSB001 predators on RSB005 prey, n = 13 for RSB001 predators on RSA075 prey, RSB005 predators on RSB005 prey, and n = 10 for each of the other conditions.

Therefore, between strain competition is a key element defining aggregate composition with *P. pacificus* preferentially grouping with their own kin and close relatives and not tolerating the presence of more distantly related strains.

As aggregation behaviors were disrupted between more distantly related strains but not close relatives, we next investigated if this may be due to aggressive predatory behaviors between strains. *P. pacificus* attacks other nematodes including other *P. pacificus* strains which can result in the killing of these larvae; however, the thicker cuticle of the adults usually prevents fatal interactions and instead induces an escape response (*19*).

Additionally, the kin-recognition system prevents any aggressive interactions between a strain’s own progeny as well as their close relatives (*21*, *22*). Therefore, we utilized previously established assays (*16*) to determine the predation outcome between our selected strains. We observed no killing of self-progeny in all strains as expected and evidence of only very limited predatory interactions between the closely related RSB001 and RSB005. However, predation between either RSB001 or RSB005 with larvae of RSA075 resulted in a highly aggressive response with large numbers of larvae killed. Additionally similar severe killing was observed in the reciprocal assays utilizing RSA075 predators with prey from either RSB001 or RSB005 (Fig. 2E). Therefore, between strain aggregation correlates with their kin-recognition and associated predatory interactions. Thus, closely related strains such as RSB001 and RSB005 perceive one another as kin, do not attack and are capable of aggregating together. Conversely, more distantly related strains are regarded as non-kin, aggressively predate one another and aggregate formation is disrupted.

### Phenotypic plasticity and predatory interactions enforce kin aggregation

We next investigated the significance of the non-kin associated aggressive interactions on aggregation behaviors in more detail by removing the ability of these strains to predate one another. One mechanism to suppress the predatory behaviors in *P. pacificus* is through manipulating their phenotypically plastic mouth as during development *P. pacificus* form one of two distinct mouth morphs which are also associated with specific feeding behaviors (*16*). This irreversible developmental process results in the formation of either the stenostomatous (St) mouth type which is strictly microbivorous and is characterized by a narrower mouth opening and a single small dorsal tooth. Alternatively, animals can develop the eurystomatous (Eu) morph which promotes an omnivorous diet including predatory behaviors and is defined by a wider, shallower mouth structure, an enlarged dorsal tooth and an additional sub-ventral tooth (*46*) (Fig. 3A). The majority of *P. pacificus* strains exhibit an Eu bias under most conditions assessed which suggests a propensity for these strains to aggressively compete with other nematodes in their environment via predatory interactions (*22*, *47*). Importantly, the *P. pacificus* mouth morph fate is influenced by a panoply of well characterized genetic and environmental factors providing a mechanism to manipulate the mouth form and subsequently the *P. pacificus* predatory behavior (*47–51*). Therefore, as RSB001, RSB005 and RSA075 are all highly Eu we utilized CRISPR/Cas9 (*43*) to generate mutations in the nuclear hormone receptor *Ppa-nhr-40* which is required for the Eu mouth morph fate (*49*). Mutations in both *Ppa-nhr-40* ^RSB001^ and *Ppa-nhr-40* ^RSA075^ resulted in 100% of animals forming the St mouth form which also succeeded in abolishing their predatory behavior (Fig. 3B and 3C and Fig. S3A and S3B).

**Figure 3.**
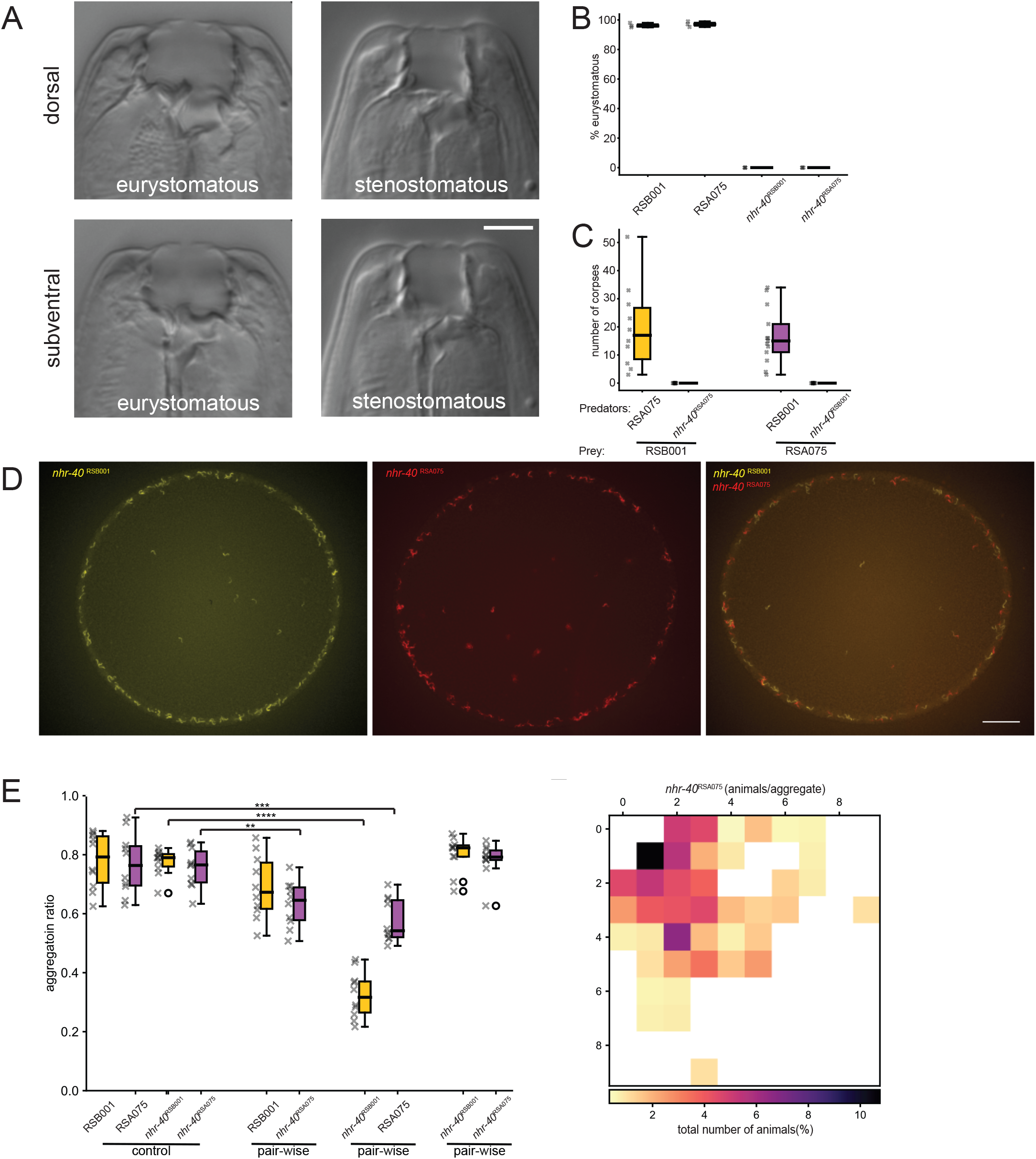
Mouth form plasticity and associated predation behaviors influence aggregation. (A) *P. pacificus* mouth dimorphism. The eurystomatous mouth has a wide buccal cavity with two teeth and is omnivorous feeding on both bacteria and other nematodes while the stenostomatous mouth is narrow with a single tooth and microbivorous. Scale bar = 5 µm (B) Mutations in *Ppa-nhr-40* in both RSB001 and RSA075 change the mouth form frequency from highly Eu to 100% St. (C) Killing assays comparing the ability of RSA075 and *Ppa-nhr-40* ^RSA075^ to predate upon RSB001 prey and the reciprocal assays of RSB001 and *Ppa-nhr-40* ^RSB001^ to predate upon RSA075 prey. Mutations in *Ppa-nhr-40* ^RSB001^ and *Ppa-nhr-40* ^RSA075^ remove the ability of these strains to predate one another. n = 10 per condition. (D) Representative images of the aggregation phenotypes of the *P. pacificus Ppa- nhr-40* ^RSB001^ and *Ppa-nhr-40* ^RSA075^ controls as well as mixed cultures of *Ppa-nhr-40* ^RSB001^ and *Ppa-nhr-40* ^RSA075^ which aggregate together. Scale bar = 2000 µm. (E) Quantification of aggregation ratios in *P. pacificus* cultures including of RSB001, RSA075, *Ppa-nhr-40* ^RSB001^, and *Ppa-nhr-40* ^RSA075^ controls as well as all pairwise permutations. n = 10 replicates. p<0.001 for RSA075 with *nhr-40* ^RSB001^, p<0.0001 for *nhr-40* ^RSB001^ with RSA075, and p<0.001 for *nhr-40* ^RSA075^ with RSB001. (F) Heatmap of pairwise interactions between *Ppa- nhr-40* ^RSB001^ and *Ppa-nhr-40* ^RSA075^ revealing the proportion of each strain in an aggregate. Mixed *Ppa-nhr-40* ^RSB001^ and *Ppa-nhr-40* ^RSA075^ mutant cultures show higher frequencies of between strain aggregates than RSB001 and RSA075 wild type strains.

With the *nhr-40* ^RSB001^ and *nhr-40* ^RSA075^ non-predatory variants established, we next explored the influence of their predatory behaviors on group selectivity and aggregate formation. In pairwise aggregation assays between the two mutant strains, *nhr-40* ^RSB001^ and *nhr-40* ^RSA075^, we observed between strain aggregation occurring abundantly confirming non-kin associated predation is a significant factor influencing the collective dynamics in *P. pacificus*. However, to investigate this further, we also conducted assays between *nhr-40* mutants together with their rival strain Eu morph. With only one strain capable of predatory behaviors in these assays we predicted the Eu strain would dominate and form aggregates at the expense of its *nhr-40* mutant opponent. Indeed, in assays of RSA075 together with *nhr-40* ^RSB001^ the highly St mutant strain was significantly displaced from grouping and aggregates were instead dominated by the predatory Eu RSA075 animals (Fig. 3D and 3E). However, in assays between RSB001 and *nhr-40* ^RSA075^, both strains unexpectedly still formed aggregates. As such, the capability of RSB001 to attack *nhr-40* ^RSA075^ and the inability of RSA075 to reciprocate with any aggression of its own did not disrupt between strain aggregation (Fig. 3D and E). Therefore, RSA075 aggregation is more robust and difficult to displace than that observed in RSB001 likely indicating the existence of additional factors which contribute to the induction and maintenance of aggregate formation in *P. pacificus*.

Thus, phenotypic plasticity and its associated predatory interactions are key components influencing aggregation behaviors, however, other yet to be determined elements also contribute to the ability of a strain to form and sustain aggregates.

### Mutations in *self-1* are insufficient to disrupt aggregation

With aggregate formation observed between animals of the same genotype as well as between kin but not more distantly related strains, we next explored the importance of the *P. pacificus* molecular kin-recognition components for these interactions. The only molecular component involved in this process that has been described so far is *self-1* which encodes for a small peptide containing a hypervariable C-terminus that is thought to be important for generating between strain specificity. Mutations in *self-1* cause a mild kin-recognition defect resulting in predators erroneously killing their own offspring (*21*). *self-1* has been previously identified in all of our three selected assay strains (*22*) (Fig. S4). As RSB001 has previously formed the basis of numerous molecular studies we focused on this strain (*25*, *27*, *43*). We generated a putative *self-1* null mutant (*self-1.1*) in RSB001, however surprisingly no kin-recognition defect was observed (Fig. 4A and B). Therefore, we analyzed the RSB001 genome further and identified a further potential *self-1* paralogue which shares 72.9% sequence identity at the amino acid level outside of the hypervariable domain (Fig. 4A). This additional copy was designated *self-1.2* and a subsequent CRISPR/Cas9 induced putative null mutation successfully phenocopied the modest kin-defective phenotype previously described in *self-1* mutants in other strains (Fig. 4A and B) (*21*). Additionally, a *self-1.1; self-1.2* double mutant revealed a stronger kin-killing defect than the *self-1.2* single mutant alone

**Figure 4.**
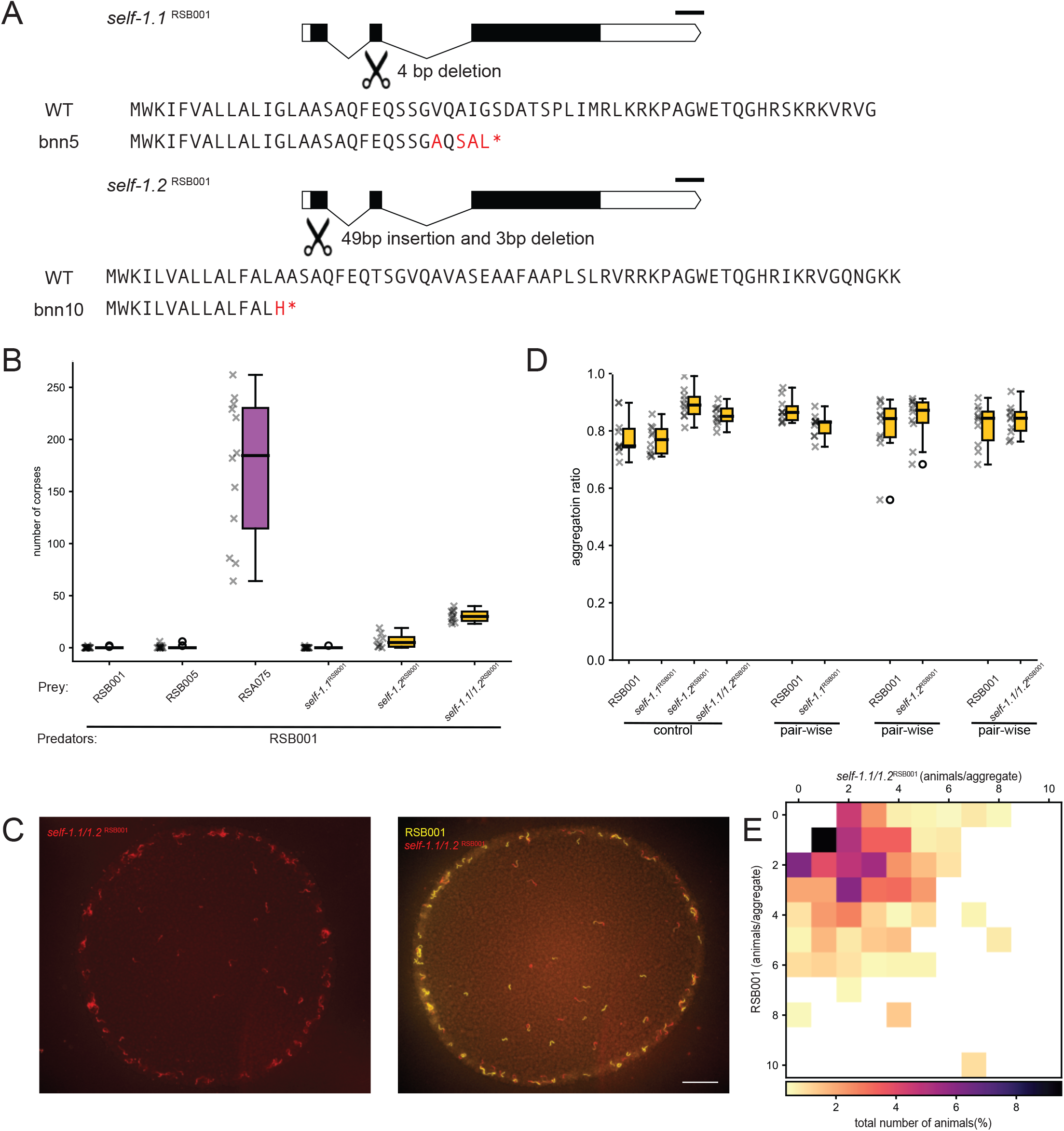
The kin-recognition component SELF-1 is dispensable for aggregate formation. (A) Predicted gene structure of *self-1.1* and self-1.2 in strain RSB001. CRISPR/Cas9 target sites are highlighted across both genes. Mutations generated resulted in severely truncated proteins and putative null mutants. Predicted wildtype protein sequences as well as associated mutations are shown. Scale bar = 100 bp. (B) Quantification of predation assays revealing the modest kin-killing phenotype caused by mutations in *self-1.* n = 21 for RSB001 prey, n = 13 for RSB005 prey, n = 12 for RSA075 prey and n = 10 for *self-1.1*^RSB001^ prey, *self-1.2*^RSB001^ prey and *self-1.1; self-1.2*^RSB001^ prey. (C) Representative images of the aggregation phenotypes of the *self-1.1; self-1.2*^RSB001^ mutant as well as mixed cultures of *self-1.1; self-1.2*^RSB001^ with RSB001. Scale bar = 2000 µm. (D) Quantification of aggregation ratios of *P. pacificus* cultures including of RSB001 and *self-1* mutants alone as well as mixed cultures of RSB001 together with *self-1* mutants. Mutations in *self-1.1; self-1.2*^RSB001^ are not sufficient to disrupt aggregation. n = 10 per condition. (E) Heatmap of pairwise interactions between of RSB001 and *self-1.1; self-1.2*^RSB001^ revealing the proportion of each strain in an aggregate. Mixed RSB001 and *self-1.1; self-1.2*^RSB001^ cultures aggregate together.

(Fig. 4B). Therefore, we next assessed if the kin-recognition defect associated with these *self-1* mutations was sufficient to also disrupt aggregation behaviors. In aggregation assays consisting of either mutants alone, or assays containing an equal amount of RSB001 together with *self-1* mutants, aggregation was maintained at wild type control levels despite the kin-recognition defect (Fig. 4B to D). Therefore, the mild kin-recognition defect caused by these mutations is insufficient to disrupt aggregation. With recent studies demonstrating that additional as yet unidentified components must also contribute to the kin-recognition signal (*22*), we predict that mutations in these other elements may result in stronger kin-recognition defects which could be sufficient to also influence and disrupt aggregation behavior.

### *P. pacificus* territorial biting prevents *C. elegans* aggregation

Interactions between con-specific *P. pacificus* likely occur frequently in their natural habitat as strains compete to occupy their preferred ecological niche. However, *P. pacificus* also comes into contact with other nematode species including *C. elegans* and its predatory ability has previously been implicated in territorial behaviors against this species (*19*, *52*).

Therefore, we next assessed conspecific aggregation interactions between diverse *C. elegans* strains as well as the impact of *P. pacificus* on the aggregation ability of these wild *C. elegans* isolates. We selected two strains of *C. elegans* which both aggregate strongly, CB4856 isolated from the Hawaiian Islands, and JU2001 from La Reunion Island (Fig. 5A). Utilizing the same fluorescent dye method employed to stain *P. pacificus* (*45*), we firstly explored pairwise aggregation assays between these con-specific *C. elegans* strains. Here, both strains successfully aggregated with one another at similar levels to with their own strain indicating the presence of another distantly related rival did not disrupt aggregation in *C. elegans* (Fig. 5A to C). Next, we challenged the *C. elegans* aggregating strains with the addition of the aggregating *P. pacificus* strain RSB001. In these mixed species assays, we observed *C. elegans* failed to aggregate, were unable to congregate at the bacterial border and instead behaved in a solitary manner. Furthermore, the aggregates were now dominated by *P. pacificus* (Fig. 6A to C). Finally, as *P. pacificus* strongly displaces *C. elegans* from aggregates, we analyzed the potency of *P. pacificus* for influencing these *C. elegans* behaviors by reducing the representation of *P. pacificus* present in mixed species pairwise assays. Here, we observed *C. elegans* aggregation behaviors were disrupted when only 17% of the assay population was *P. pacificus* (Fig. 6D and E). Additionally, under these conditions *C. elegans* aggregates were mostly only maintained when they consisted of large numbers of animals. Therefore, *C. elegans* aggregates above a density threshold may be sufficient to resist predation induced displacement and may represent an ecological relevant strategy to counter the effects of *P. pacificus* predation (Fig. 6F). Thus, a minimal *P. pacificus* presence is capable of shaping the aggregation behaviors of other species such as *C. elegans*. These interactions may therefore represent a territorial behavior conferring an ecologically relevant and advantageous state for *P. pacificus* within their shared environmental niche.

**Figure 5.**
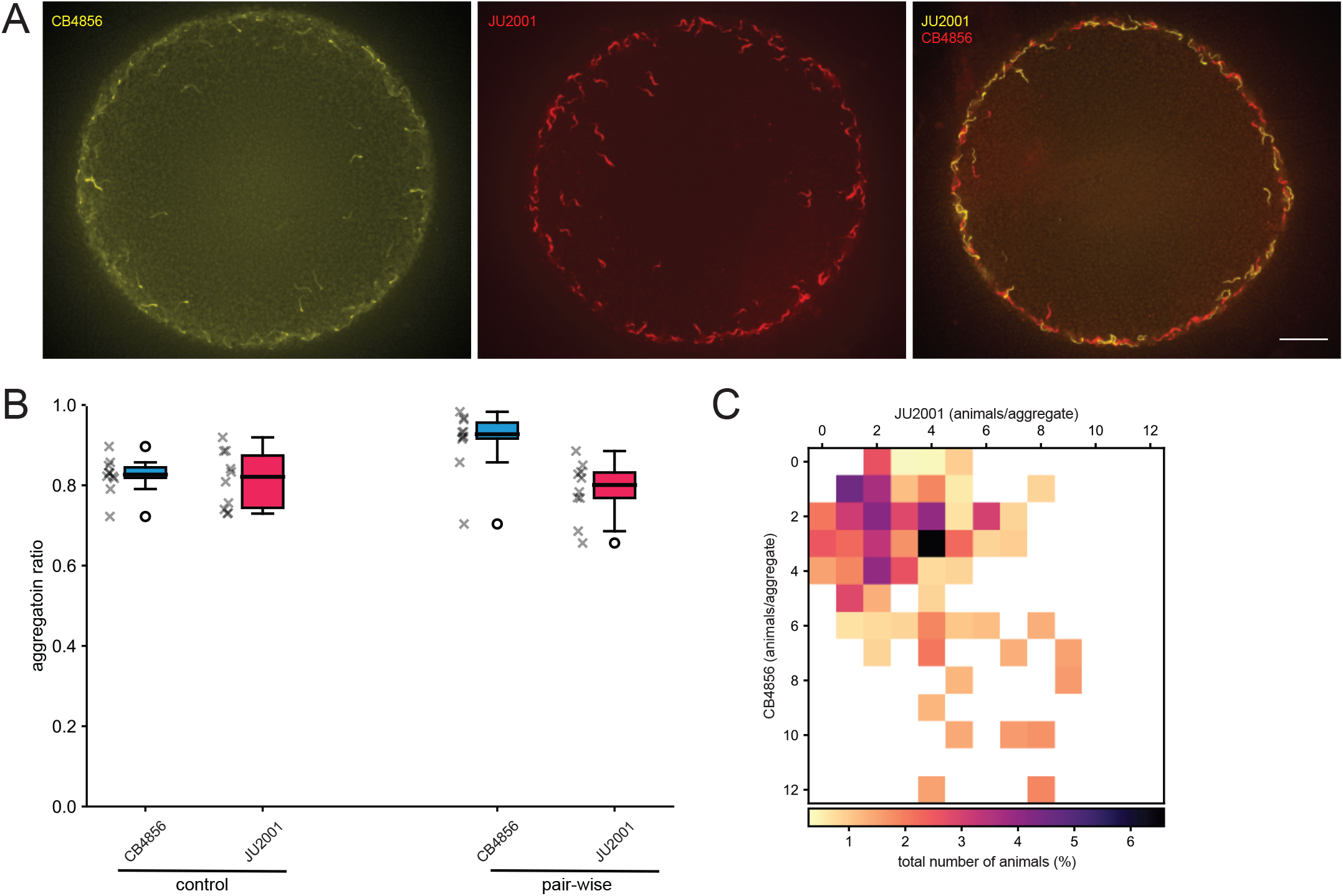
***C. elegans* pairwise interactions between aggregating strains.** (A) Representative images of the aggregation phenotypes of the *C. elegans* strains CB4856 (Hawaiian) and JU2001 (La Reunion) controls as well as mixed cultures of CB4856 and JU2001 together. Scale bar = 2000 µm. (B) Quantification of aggregation ratios of *C. elegans* cultures including of CB4856 and JU2001 alone as well as mixed cultures of CB4856 and JU2001 together. Both C. elegans strains aggregate together abundantly. n = 10 per condition. (C) Heatmap of pairwise interactions between of CB4856 and JU2001 revealing the proportion of each strain in an aggregate. Mixed CB4856 and JU2001 cultures form frequent aggregates together.

**Figure 6.**
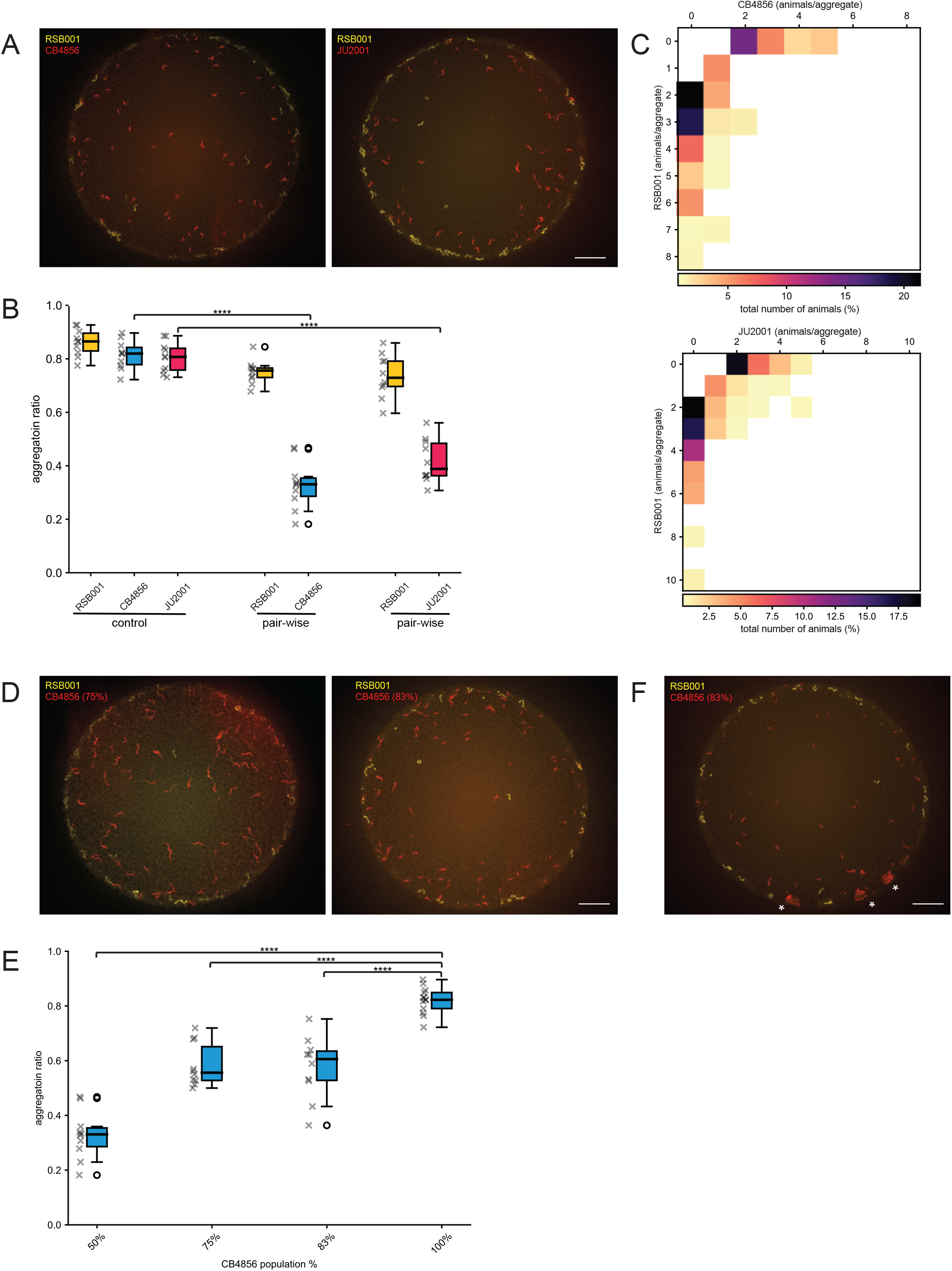
***P. pacificus* interferes with *C. elegans* aggregation behaviors.** (A) Representative images of the aggregation phenotypes with *P. pacificus* in mixed cultures together with *C. elegans* strains CB4856 (Hawaiian) or JU2001 (La Reunion). Scale bar = 2000 µm. (B) Quantification of aggregation ratios of *P. pacificus* RSB001 and *C. elegans* CB4856 and JU2001 controls as well as mixed cultures of RSB001 together with CB4856 or JU2001. RSB001 disrupts the aggregation ability of both *C. elegans* strains. n = 10 per condition. p<0.0001 for CB4856 and JU2001 with RSB001. (C) Heatmap of pairwise interactions between *P. pacificus* RSB001 and *C. elegans* CB4856 or JU2001 revealing the proportion of each strain in an aggregate. Monotypic aggregates are more prevalent when *C. elegans* is paired with *P. pacificus* RSB001 which also dominates and reduces aggregate formation in the *C. elegans* strains. (D) Representative images of the aggregation phenotypes with a reduced ratio of *P. pacificus* to *C. elegans* CB4856 in mixed cultures assays. Scale bar = 2000 µm. (E) Quantification of aggregation assays with a reduced ratio of *P. pacificus* RSB001 to *C. elegans* CB4856. Even a minimal *P. pacificus* presence is capable of shaping the aggregation behaviors of *C. elegans.* n = 13 for 100% CB4856 and n = 10 for each of the other conditions. p<0.0001. (F) Representative aggregation assay with a reduced ratio of *P. pacificus* RSB001 to *C. elegans* CB4856. Occasionally *C. elegans* aggregates were established consisting of large numbers of *C. elegans* (marked *) which may be sufficient to resist predation induced displacement. Scale bar = 2000 µm.

## Discussion

Group behavior in nematodes including aggregate formation emerge from local interactions between individuals (*32*). In *P. pacificus* however, an additional layer of complexity is evident through its predatory capacity which functions to broaden potential food opportunities and as a mechanism to remove competitors from their local environment (*18–22*) as well as its kin-recognition ability which prevents attacks on close relatives (*21*, *22*). In our work, we show these elements combine to also impact their collective behaviors as aggregation is favored between kin while predatory attacks between distantly related con- specifics result in the displacement of their rivals and the induction of less preferable behavioral strategies. Moreover, in pairwise assays between distantly related strains, we also observed that one strain is able to dominate these interactions resulting in monotypic aggregates. Importantly, the exclusion of potential competitors from optimal kin groupings is not unique to *P. pacificus* and has been observed in many organisms. This includes in slime molds which aggregate together in fruiting bodies under stressful conditions. This behavior depends on the between strains compatibility of the cell surface *csA* gene which ultimately facilitates admission to the fruiting body (*5*, *53*). Additionally, in social yeast species, robust cell adhesion molecules promote efficient homogenous aggregate and biofilm formation while weak adhesive forces between non-kin result in their exclusion (*2*, *54*). However, these mechanisms of selectivity and omission depend on preventing aggregations between non-kin which differs from our observations in *P. pacificus* whereby predation provides an active mechanism to displace competitors.

Subsequently, we also demonstrated that the aggressive displacement of potential competitors from aggregates is at least partially dependent on the predatory Eu mouth form. This is the most prevalent morph found in wild *P. pacificus* isolates including those from the high altitude adapted clade utilized in these studies (*22*, *47*). As all strains of *P. pacificus* are capable of also developing the non-predatory St morph, aggregation behaviors are therefore theoretically possible even between highly divergent strains which may prove to be beneficial under certain environmental conditions. However, despite the influence of the mouth form and its associated predatory behaviors on aggregate formation, our interaction experiments with St morph animals demonstrated that other factors must also determine the robustness of aggregates. Specifically, RSA075 appears to form more durable aggregates than RSB001. Perhaps, RSA075 is less responsive to predatory attacks than RSB001 or alternatively, it may form a stronger kin-association. In addition to the role of *P. pacificus* predation, previous studies have identified several genetic components that are necessary for *P. pacificus* aggregation. In particular, mutations in the IFT system cause a solitary strain of *P. pacificus* to aggregate (*25–27*, *36*). Therefore, in the future, it will be important to assess the impact of these mutations on con-specific aggregation behaviors which may aid in establishing more of the sensory mechanisms involved however, due to their substantial influence on mouth morph fate, these experiments will not be straightforward.

While predation and kin-recognition appear to play an integral role in aggregation behaviors in *P. pacificus*, neither behavior has been observed in *C. elegans*. Accordingly, we observed aggregation between distantly related social strains of *C. elegans* with no apparent effect on their behaviors. This is consistent with previous studies in which a mixed population of *C. elegans* including an aggregating strain and an evolutionary distinct solitary strain, each maintained their specific behavioral phenotypes when grown together (*34*). However, in mixed assays between aggregating *C. elegans* and *P. pacificus* strains, aggregation was hindered in *C. elegans* while maintained in *P. pacificus*. Furthermore, the presence of relatively few *P. pacificus* predators proved sufficient to modulate the aggregation behavior in *C. elegans* suggesting a minimal amount of *P. pacificus* animals are capable of altering the group dynamics of a much larger competing nematode population. As *P. pacificus* develops slower and have a smaller brood size than *C. elegans* (*55*, *56*), the ability to alter the behavior and outcompete other nematodes such as *C. elegans* may therefore be highly beneficial. Furthermore, this may represent an additional facete to the previously characterized *P. pacificus* territorial behaviors which have been shown to disrupt *C. elegans* ability to approach and enter shared bacterial lawns (*19*).

With our work revealing a previously unknown influence for kin-recognition and its associated predation on collective behaviors in *P. pacificus,* questions remain regarding what other behaviors may also be affected by these abilities. For example, recent work in *C. elegans* has demonstrated that their swimming gait is influenced by the presence of other nearby animals (*57*). Additionally, more diverse collective behaviors including swarming have been described in *C. elegans* (*31*, *32*) and groups of both *C. elegans* and *P. pacificus* are capable of forming large 3D tower like structures thought to aid in their dispersal to new environments (*58*, *59*). Therefore, how predation and kin-recognition may influence these other collective behaviors remains unknown and will form the basis of future studies. Thus, kin-recognition abilities in *P. pacificus* facilitate a diverse behavioral repertoire of interactions. This includes the territorial removal of competitors via predatory events, as well as between kin-aggregation in which individuals nepotistically favor grouping with relatives as rivals take on less preferable behavioral strategies. Furthermore, with the wealth of molecular tools available, *P. pacificus* offers a powerful means for understanding the genetic and neural mechanisms behind these kin-recognition mediated interactions and their evolution.

## Materials and Methods

### Nematode Husbandry

All nematodes used were maintained on standard NGM plates on a diet of *Escherichia coli* OP50. All strains used in this study can be found in the supplementary table S1.

### Aggregation assays

Aggregation assays were utilized to quantify aggregation behaviors between different strains. Assay plates were prepared two days prior to the experiment. 6 cm NGM plates were seeded with 70 µl of OP50 bacteria and incubated at room temperature. All animals were maintained on NGM plates seeded with OP50 bacteria until freshly starved, resulting in an abundance of young larvae. These plates were washed with M9 and passed through two 20 µm filters to isolate pure cultures of larvae. They were centrifuged and transferred onto new NGM plates seeded with OP50 bacteria and incubated at 20 °C for three days until they become young adults. They were then washed with M9 and transferred into 1.5 ml tubes, containing 50 µM of dye (CellTracker™ Green BODIPY™ Dye or CellTracker™ Orange CMRA Dye), and incubated on a rotator in dark at 20 °C for 3 h. The worms were then washed with M9 four times and transferred into an unseeded NGM plate. As control 120 worms from one strain were picked and transferred to an assay plate, outside of OP50 bacteria. For pairwise aggregation assays, 60 worms per strain from two strains were transferred to an assay plate. The plates were incubated at 20 °C in dark for 3 h, then imaged using a fluorescent microscope ZEISS Axio Zoom.V16 with ZEN (blue edition) software. Images were analyzed manually with Fiji. It was considered an aggregate when two or more animals were in contact. The staining with CellTracker™ dyes method was adapted from Werner et al (*45*) and pairwise aggregation assay was adapted from Moreno et al (*25*).

### Off food aggregation assay

Off food aggregation assays were utilized to investigate aggregation behaviors of nematodes in an environment without bacterial food (OP50). Animals are well-fed and are washed prior to the experiment. 360 worms of the same strain were transferred on a 3.5 cm NGM plate.

The plates were incubated at 20 °C for 1 h. Images were taken using brightfield on microscope ZEISS Axio Zoom.V16 with ZEN (blue edition) software.

### CRISPR/Cas9 induced mutations

Mutations were induced in candidate genes via CRISPR/Cas9. Gene specific crRNA and universal trans-activating CRISPR RNA (tracrRNA) was purchased from Integrated DNA Technologies and 5 μL of each 100 μM stock mixed and denatured at 95°C for 5 min before cooling at room temperature to anneal. Cas9 (Integrated DNA Technologies) was added to the hybridized product and incubated at room temperature for 5 mins. This was subsequently diluted with TE buffer to a final concentration of 18.1 μM for the sgRNA and 2.5 μM Cas9. This was injected into the germline of the required strain *P. pacificus*. Eggs from injected P0s were recovered up to 16 h post injection. After hatching and 2 days’ growth these F1 were segregated onto individual plates until they had also developed sufficiently and egg laying had been initiated. The genotype of the F1 animals were subsequently analyzed via Sanger sequencing and mutations identified before re-isolation in homozygosis. sgRNAs and associated primers utilized in this study can be found in Table S2.

### Mouth-form

Mouth form phenotyping was achieved by observation of the nematode buccal cavity using ZEISS Axio Zoom.V16 microscope with morph identities based on previous described species characteristics (*47*). Final mouth-form frequencies are the mean of 3 independent replicates, each assaying 100 animals.

### Predation and Kin-recognition Assays

Corpse assays facilitated rapid quantification of predatory behaviors between different strains. Prey were maintained on NGM plates seeded with OP50 bacteria until freshly starved, resulting in an abundance of young larvae. These plates were washed with M9 and passed through two 20 µm filters to isolate pure cultures of larvae. They were subsequently centrifuged before being deposited on to an unseeded assay plate by pipetting 1.5 µl of *P. pacificus* or 1.0 µl of *C. elegans* larval pellet on to a 6 cm NGM unseeded plate. Five predatory nematodes were screened for the appropriate mouth morph and added to assay plates for prey assays. They were then permitted to feed on the prey for 2 h and the plate was subsequently screened for the presence of corpses. When utilizing *self-1* mutants as prey for kin-recognition assays, the predators were placed on assays and left for 20 h before the plate was subsequently screened for the presence of corpses.

### Statistical Analysis

Welch’s t-test was performed using SciPy library. Box plot represents the first quartile (Q1) to the third quartile (Q3) of the data with a line at the median (Q2). Q1, Q2, and Q3 are the values which lie at 25%, 50% and 75% of the data points. Whiskers represent the range in which most values are found, whereas values outside this range are presented as outliers. The statistical tests were always performed against the control condition of the same strain. non-significant (ns), p-value ≤ 0.05 (*), p-value ≤ 0.01 (**), p-value ≤ 0.001 (***), p-value ≤ 0.0001 (****).

## Acknowledgements

We would like to thank Marianne Roca and Monika Scholz for discussion and critical reading of the manuscript (MPI Neurobiology of Behavior – caesar, Bonn). Additionally, we wish to thank the Sommer lab for *P. pacificus* strains, Christian Rödelsperger for phylogenetic tree data and bioinformatic discussions (MPI for Biology, Tübingen), Marie-Anne Felix and Aurélien Richaud for *C. elegans* strain JU2001 (IBENS, Paris) and Bogdan Sieriebriennikov (New York University) for mouth form images. Finally, some strains were provided by the CGC, which is funded by NIH Office of Research Infrastructure Programs (P40 OD010440).

## Author Contributions

Conceptualization: FH, JWL

Methodology FH, JWL

Investigation FH

Visualization FH

Supervision JWL

Writing – original draft: JWL

Writing – review and editing FH, JWL

## Data and Materials Availability

All data are available in the main text or the supplementary materials. All data and nematode strains are available upon request.

