## Supplemental Files for "Kin-recognition shapes collective behaviors in the cannibalistic nematode *Pristionchus pacificus*"

#### Supplementary Text

**Fig. S1.** The solitary *C. elegans* strain N2 and social CB4856 both do no aggregate in the absence of bacteria. The *P. pacificus* PS312 solitary strain also does not aggregate in the absence of food however the social RSB001 continues to aggregate. Scale bar = 2000 μm.

**Fig. S2.** (A) and (B) Both RSB005 and RSA075 form frequent aggregates under standard laboratory conditions which is not affected by CellTracker™ Green BODIPY™ Dye or CellTracker™ Orange CMRA Dye staining. Scale bar = 2000 μm.

**Fig. S3.** (A) Gene structure and CRISPR target site for *Ppa-nhr-40* mutations in both RSB001 and RSA075. Mutations were successfully generated in both strains. Scale bar = 1kb. (B) Mutations result in a frame shift in both RSB001 and RSA075 strains which leads to a putative premature stop codon and a truncated protein.

**Fig. S4.** *self-1* is present in all three strains used in the study. This includes variable copy numbers and in RSA075 also distinct hypervariable regions.

**Table S1.**
List of all strains used and alleles associated with the mutations.

**Table S2.**
List of primers and CRISPR/Cas9 associated sequences for generating mutants.

**Movie S1.**
Movie showing pairwise assay between two competing strains. RSA075 is stained yellow and RSB001 is stained in red.

### Supplementary Fig. 1

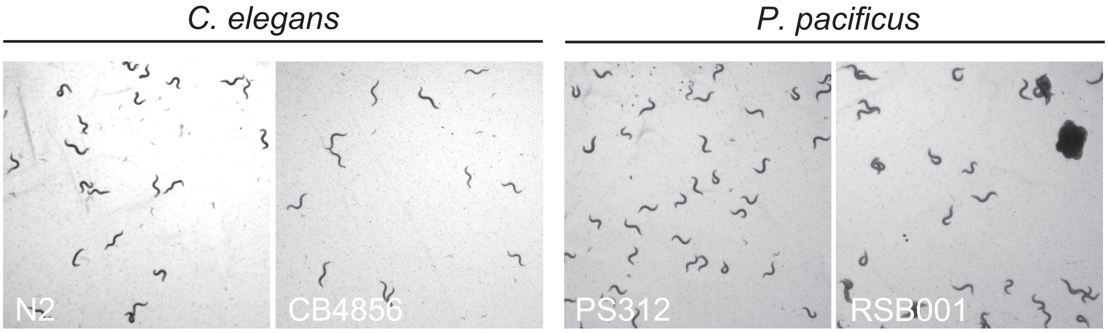

### Supplementary Fig. 2

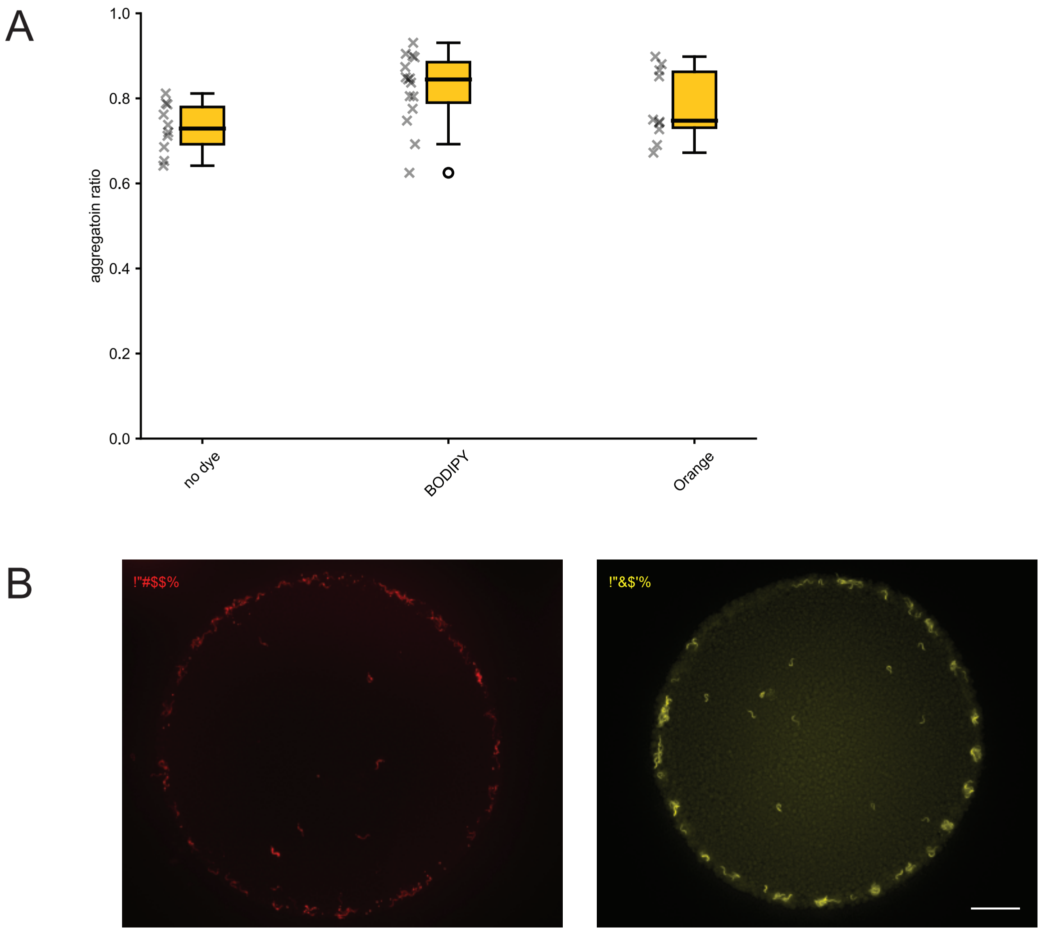

### Supplementary Figure 3

## A

*Ppa-nhr-40*

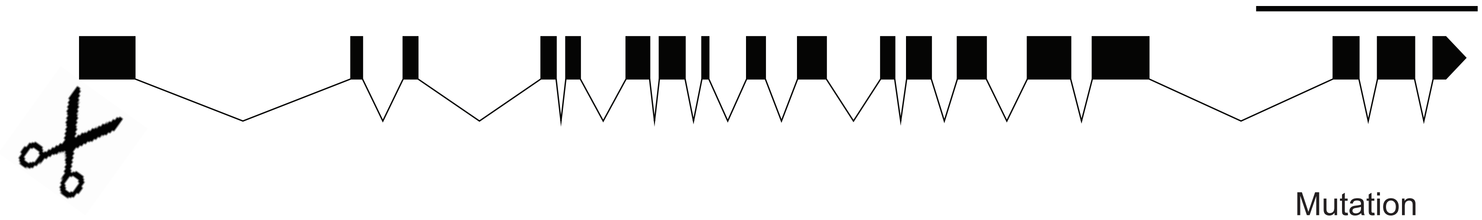

| | | Mutation |
|---|---|---|
| *Ppa-nhr-40* | GAAGATTCCGGAGGG-AACCCTCTGTGTCGT | Wild type |
| *Ppa-nhr-40* RSB001 *(bnn38)* | GAAGATTCCGGAGGGGAACCCTCTGTGTCGT | 1bp insertion |
| *Ppa-nhr-40* RSA075 *(bnn40)* | GAAGATTCCGGAG-----CCCTCTGTGTCGT | 4bp deletion |

## B

*Ppa-nhr-40*

MEIYGRCDYTTTHRVEIYTRREKIPEGTLCVVCDDSASGIHYSVASCNGCKTFFRRALVN
KQTFTCQFEGKCLVGKNVRCVCRSCRLKKCFEQGMDPKAIQHDRDKIRYTKVLKREKEAL
KAKKEAERESMMMKVKEEIGSPGGLECCDIPSTSKGNPFSFLSPMEMILSRFLSNEPSND
LDSNLRELMRIEKKVVEVRNAYRYDDQISTIWNMYIGSRAMLSEDDWLSATTQQKPMLFS
LSERQEPAQVSEPPKTSPYRCARITPWSLREWFQRDLTLMMEWVKLIPGINDLITSDKVI
LQKNFALTFAVYQLTFYTMDTTVLSGDDFAPSELSSLEERLKSLKRRSDSRSTPPPPAPC
LSSTINNELMNLANSLYKQPLAKRIKDEILEDEPTCSAYLRKLTEQASLLATSSPSTSSF
VNSPIDSAFLMKNGLTAVSSFSEGINQFPSNMPPLPPTMSTGLPGALSNCLASSLQAASK
NMTSLPTTLPTFPSGFFSALHNSPASLGFPPSPLLASLTASPLITSSPLMAQSPLMAQSP
LMASLTMPSHSLPSTFPNFLPPPVPAHLALPPPSLPTLLIPLLPQPRPLPVNQLPPAPVP
IREKTPDIDVETVSMDGRSEGRGSRAIDTTDSRDEVLFRVPASLPRSISVTSSSDVIKPF
NLMSMRSPDDPSRTVKMDSILEKMSVSPAEQMTIPSTIVSSVMSTSTSSNNIKDEPESPE
EMTSSNSSSVIKNTGDVTEITKEPKVHKPQAENRTMFEDEETPCDPTPEAQIANRINYPD
GTFIERDKERPFNDELYGLLIDGIWKIFRRENVDQETFVLFKMMSFFNTELTHKGDHKLS
EDGVKYVEKMRQKMYTQLLLHLQKTKGKDGLRIFSSLLLMGSTIARVRNALRKMFTMTSI
FVPSNDLVDQLILRDNDERVPSPSAFSIYHPVY*

*Ppa-nhr-40* RSB001 *(bnn38)*

MEIYGRCDYTTTHRVEIYTSGQNSDKTLILMLFPGREKIPEGNPLCRL*

*Ppa-nhr-40* RSA075 *(bnn40)*

MEIYGRCDYTTTHRVEIYTSGQNSDKTLILMLFPGREKIPEPSVSSVMIRPLAFTILSLRAM
DAKPSSEELS*

### Supplementary Figure 4

| | | hypervariable domain |
|---|---|---|
| SELF-1$^{RSB001}$ | MWKILVALLALIGLAASAQFEQSSGVQAIGSDATSPLIMRLKRKPAGWETQGHRSKR | KVRVG--- |
| SELF-1$^{RSB005}$ | MWKILVALLALIGLAASAQFEQSSGVQAIGSDATSPLIMRLKRKPAGWETQGHRSKR | KVRVG--- |
| SELF-1.1$^{RSA075}$ | MWKILVALLALIGLAASAQFEQSSGVQAIGSDATSPLIMRLKRKPAGWETQGHRSKR | V------- |
| SELF-1.2$^{RSA075}$ | MWKILVALLALIGLAASAQFEQSSGVQAIGSDATSPLIMRLKRKPAGWETQGHRSKR | I------- |

Supplementary Table S1

| **Name** | **Strain** | **Alleles** | **Species** |
|---|---|---|---|
| RSB001 | RSB001 | Wild Type | *P. pacificus* |
| RSB005 | RSB005 | Wild Type | *P. pacificus* |
| RSA075 | RSA075 | Wild Type | *P. pacificus* |
| *self-1.1* | JWL8 | *self-1.1 (bnn5)* | *P. pacificus* |
| *self-1.2* | JWL15 | *self-1.2 (bnn10)* | *P. pacificus* |
| *self-1.1; self-1.2* | JWL90 | *self-1.1 (bnn57); self-1.2(bnn10)* | *P. pacificus* |
| *nhr-40* (RSB001) | JWL52 | *nhr-40 (bnn38)* | *P. pacificus* |
| *nhr-40* (RSA075) | JWL54 | *nhr-40 (bnn40)* | *P. pacificus* |
| CB4856 | CB4856 | Wild Type | *C. elegans* |
| JU2001 | JU2001 | Wild Type | *C. elegans* |

Supplementary Table S2

| Gene name | F Primer sequence | R Primer sequence | sgRNA (PAM) |
|---|---|---|---|
| self-1.1 | CGCTGTGCTGACAGTATCAAGATG | GTTTTGCACACCCCTGTTGCATAG | CGAGCAGTCTTCCGGGGTTC(AGG) |
| self-1.2 | GAATAAGTGAAGAAGGGCGG | GTTTACGCAATTCGGGTCAC | TCACTGAGCGGAAGCAGCGA(GGG) |
| nhr-40 (RSB001) | GCGTTCTTTGAAGAGACCTC | GAGTATCCCAAGGGAATCCTAC | ACGACACAGAGGGTTCCCTC(CGG) |
| nhr-40 (RSA075) | GCGTTCTTTGAAGAGACCTC | GGTACTTCCTCTTTCTCTCACG | ACGACACAGAGGGTTCCCTC(CGG) |
